# Insights into the *foraging* Gene’s Influence on Mating Investments of Male *Drosophila*

**DOI:** 10.1101/2024.07.14.603413

**Authors:** Wengjing Li, Yongwen Huang, Woo Jae Kim

## Abstract

The *foraging* gene is a key genetic factor that modulates social behavior in insects, primarily by governing the trade-off between individual foraging and group-related activities. It has been associated with various behaviors associated with food search and resource exploitation, thereby playing a crucial role in determining the efficiency of foraging and the overall success of the collective. In this study, we investigate the critical role of the *foraging* gene in mediating complex interval timing behaviors, particularly mating duration, in the fruit fly *Drosophila melanogaster*. By examining two distinct variant phenotypes, rover and sitter, we observe specific deficiencies in longer (LMD) and shorter mating duration (SMD) behaviors, respectively, suggesting the gene’s crucial influence on these interval timing mechanisms. Utilizing single-cell RNA sequencing and knockdown experiments, we identify the gene’s significant expression in key neurons involved in learning and memory. However, its impact on mating duration is not observed in these brain regions. Instead, our data reveal the gene’s crucial role in specific neurons expressing Pdfr, a critical regulator of circadian rhythms. Furthermore, the study uncovers sexually dimorphic expression patterns in the brain and highlights the necessity of the gene’s dosage in specific cell populations within the ellipsoid body for normal mating duration. These findings underscore the *foraging* gene’s pivotal role in mediating complex interval timing behaviors in *Drosophila*, providing valuable insights into the intricate interplay between genetics, environment, and behavior. This research contributes to a deeper understanding of the genetic and neural mechanisms underlying complex interval timing behaviors, with broader implications for unraveling the function of *foraging* gene.

## INTRODUCTION

The *foraging* gene (*for*) has emerged as a pivotal factor in the regulation of behavioral plasticity and decision-making processes across various species, including the fruit fly *Drosophila melanogaster* ^1,2^ and honeybee ^3^. This gene plays a crucial role in shaping an organism’s foraging strategies by influencing traits such as olfactory learning and memory^4–10^. The study of the *foraging* gene has provided valuable insights into the complex interplay between genetics and behavior, highlighting its significance in understanding the adaptive behaviors of organisms in response to their environment ^3,6^.

Interval timing behaviors, the ability to measure and respond to the passage of time, are integral to various aspects of an organism’s life, including foraging, mating, and social interactions ^11,12^. These behaviors are thought to be mediated by neural circuits that are conserved across species^13^. In recent years, researchers have discovered a remarkable connection between interval timing behaviors and social behaviors, suggesting a shared neural and genetic basis ^14^.

The mating duration of male fruit flies, *Drosophila melanogaster*, serves as an excellent model for studying interval timing behaviors. In *Drosophila*, two notable interval timing behaviors related to mating duration have been identified: Longer-Mating-Duration (LMD), which is observed when males are in the presence of competitors and extends their mating duration ^15–17^ and Shorter-Mating-Duration (SMD), which is characterized by a reduction in mating time and is exhibited by sexually experienced males ^18,19^.

Mating duration represents a critical investment of time for male reproductive success, highlighting its significance as a model for investigating interval timing. This parameter offers a valuable opportunity to explore the evolutionary and physiological roles of genes involved in this complex behavior ^20^. From an evolutionary perspective, the ability to accurately regulate mating duration allows males to maximize their reproductive output by optimizing their investment of time and energy. This time investment strategy is crucial for competing with other males and ensuring successful fertilization of female eggs ^19^. Physiologically, mating duration is influenced by a complex interplay of neural circuits, hormonal signals, and sensory inputs. Understanding the genes and mechanisms underlying this behavior can provide valuable insights into the neural substrates of interval timing and the regulation of reproductive behaviors ^15,16,18,19,21–23^.

The mating duration of male fruit flies is highly dependent on gene-environment interactions, making it a promising candidate to be regulated by the *foraging* gene function ^15,16,19^. The *foraging* gene, known for its role in mediating behavioral plasticity and decision-making processes, is likely to influence mating duration by modulating the fly’s response to environmental cues and social context. For instance, in the presence of competitors, the *foraging* gene may upregulate mating duration (LMD) to enhance reproductive success, while in situations of sexual satiation, it may downregulate mating duration (SMD) to conserve energy and resources. By examining the expression and function of the *foraging* gene in relation to mating duration, we can gain valuable insights into the intricate interplay between genetics, behavior, and environmental factors in *Drosophila*.

In this study, we present the novel finding that genes traditionally associated with foraging behavior are also involved in interval timing, a cognitive process crucial for decision-making and behavior ^12,24^. This discovery challenges the assumption that these genes are solely dedicated to optimizing foraging strategies and suggests a broader role for these genes in regulating time perception and behavior across various contexts ^13,25^. Our findings indicate that the interplay between *foraging* genes and interval timing circuits may be crucial for animals to adapt their behavior to changing environmental conditions and maximize their overall fitness. This suggests that the evolution of interval timing and foraging behaviors might be more interconnected than previously thought ^26,27^.

Understanding the mechanisms by which *foraging* genes influence interval timing can provide valuable insights into the neural and genetic basis of time perception and decision-making. This knowledge has the potential to contribute to our understanding of various behaviors, including foraging, learning, and memory, and their adaptive significance in different ecological contexts.

## RESULTS

### Two distinct *foraging* allele, rover and sitter affect distinct interval timing behaviors

The *foraging* gene in *Drosophila* gives rise to two distinct phenotypes known as rover (*for^R^*) and sitter (*for^S^*), which exhibit natural variations in foraging behavior. Rover larvae cover larger areas within food patches and greater movement between patches than sitters. Rover larvae and adults are more active than their sitter counterparts. This behavioral variation is primarily due to allelic differences in the *foraging* gene, which is located on the second chromosome and encodes a cGMP-dependent protein kinase (PKG). Rovers, which have one copy of the *foraging* gene (*for^R^*/-), exhibit higher levels of *for*-mRNA and PKG activity compared to sitters, who are homozygous for the loss-of-function allele (*for^S^*/*for^S^*). The kinase encoded by *foraging* is a critical regulator of numerous downstream targets, leading to a range of pleiotropic effects associated with the *foraging* gene ^1,28,29^.

Moreover, the neural characteristics of these phenotypes are distinctly different. Sitter neurons display higher levels of spontaneous activity and exhibit exaggerated responses to stimulation, known as excessive evoked firing, which are not observed in rover neurons. This heightened excitability in sitter neurons is attributed to a reduction in voltage-dependent potassium (K^+^) currents and the presence of excitable synapses. Additionally, the axon terminal projections of sitter strains are altered, further highlighting the neural basis for the behavioral differences between these two foraging strategies ^29^.

The *for^R^* homozygote allele displayed deficiencies exclusively in SMD behavior (Fig. 1A), while the *for^S^* homozygote allele exhibited deficits only in LMD behavior (Fig. 1B). Strikingly, the *for^R^*/*for^S^* transheterozygote exhibited deficits in both LMD and SMD behaviors (Fig. 1C), indicating that each allele specifically affects distinct interval timing behaviors. All adult flies that are heterozygous for the control demonstrate typical LMD behavior (Fig. S1A-B).

**Figure 1.**
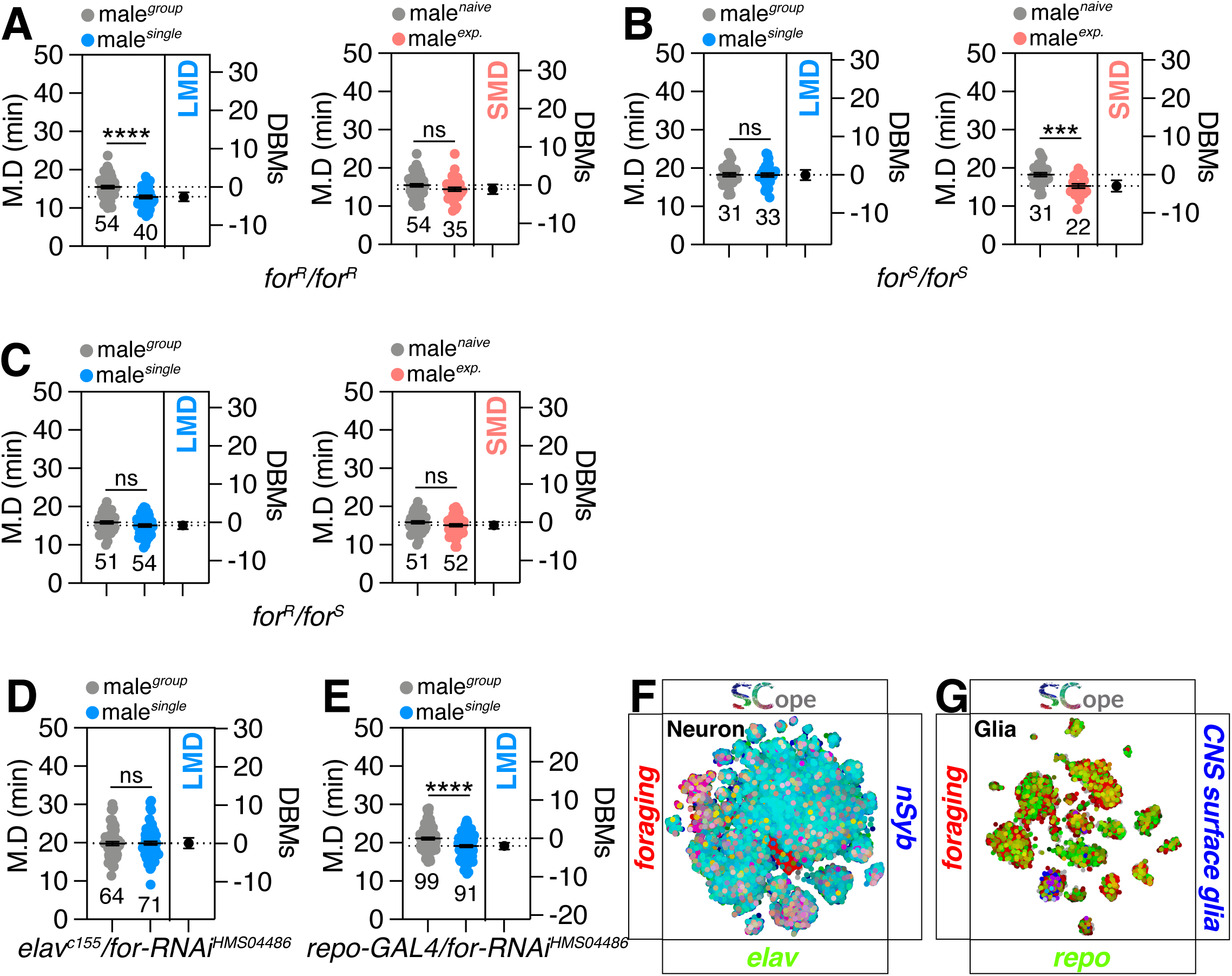
Interval behavior is regulated by two distinct *foraging* alleles. (A-C) LMD and SMD assays for *for^R^*homozygous variants, *for^S^* homozygous variants and transheterozygote *for^R^*/*for^S^*. Light grey dots in MD assays represent naïve males, blue dots represent single reared males and pink dots represent experienced males. Dot plots represent the MD (Mating Duration) of each male fly. DBMs represent difference between means. The mean value and standard error are labeled within the dot plot (black lines). Asterisks represent significant differences, as revealed by the Student’s t test (* p<0.05, ** p<0.01, *** p<0.001). The same notations for statistical significance are used in other figures. (D) LMD assays of flies expressing *elav^c155^* (neuron) driver together with *UAS-for-RNAi*. (E) LMD assays of flies expressing *repo-GAL4*(glia) driver together with *UAS-for-RNAi*. (F) Single-cell RNA sequencing (SCOPE scRNA-seq) datasets reveal cell clusters colored by expression of *foraging* (red), *nSyb/elav* (blue/green) in neurons. (G) SCOPE scRNA-seq datasets reveal cell clusters colored by expression of *foraging* (red), CNS surface glia/repo (blue/green) in glia cells.

Therefore, each *for^R^* or *for^S^* allele displays a dominant phenotype when paired with another *for^R^* or *for^S^* allele (Fig. 1C), but this dominance is lost when they are paired with a wild-type for allele (Fig. S1A-B). In molecular terms, these findings indicate that the PKG activity and regulatory mechanisms associated with each *foraging* homozygous allele is essential for disrupting LMD or SMD behaviors. Therefore, we hypothesize that an extremely high level of PKG activity specifically disrupts SMD, while an extremely low level of PKG activity specifically disrupts LMD behavior.

Given the more pronounced defects associated with the sitter allele which shows lower PKG activity ^28–30^, we have chosen to concentrate our efforts on elucidating the genes and neural circuits that underlie the influence of the *foraging* gene on LMD behavior, with the aim of mapping the precise mechanisms governing this aspect of interval timing.

### Memory circuitry for LMD behavior is not linked to the function of *foraging*

To recreate the effects of the *for^S^* allele, we employed RNA interference (RNAi)-mediated gene knockdown. This technique lowers the expression of the PKG gene, consequently reducing PKG activity in cells expressing GAL4. The neuronal knockdown of *for* using two distinctive RNAi strains disrupted LMD behavior (Fig. 1D and Fig. S1C), however glial knockdown had no effect (Fig. 1E). Utilizing the cutting-edge fly SCope single-cell RNA-sequencing data platform ^31^, we detected a significant overlap in the expression of the *foraging* gene with two key markers: *elav*, indicative of neuroblast activity, and *nSyb*, a synaptic marker, within the neuronal population (Fig. 1F). This co-expression pattern implies that the *foraging* gene likely plays a role in neuronal function. The *foraging* gene also exhibits a strong expression correlation with repo, a marker for glial cells, and is prominently expressed in specific regions of the central nervous system (CNS) surface glia (Fig. 1G). Given that RNAi and GAL4 control cross flies exhibit normal LMD behavior (Fig. S1D-E), we infer that the neuronal function of the *foraging* gene is indispensable for the generation of LMD behavior, whereas the glial function is not.

Given the *foraging* gene’s established role in learning and memory in *Drosophila* ^32^, we investigated its function in key brain regions associated with memory. We have previously demonstrated that the neural circuits responsible for the formation and retention of longer mating duration (LMD) memories are located within the R2-R4m ring neurons of the ellipsoid body (EB) ^15^. Knockdown of *for* in the mushroom body (MB) (Fig. 2A), fan-shaped body (FB) (Fig. 2B), EB (Fig. 2C), and pars intercerebrails (PI) (Fig. 2D) did not affect LMD behavior, indicating that *for* expression in these memory-related brain regions is dispensable for LMD generation. This aligns with previous findings that MB ablation via hydroxyurea feeding did not alter the locomotor activity of rover and sitter larval morphs ^33^. Our findings reveal that molecular markers for the MB (Fas2- and Mef2-positive) ^34,35^, EB (Hk- and AstA-R1-positive) ^36,37^, FB (Dh31- and ss-positive) ^38,39^, and PI (Ilp2-positive but Ilp7-negative) ^40^ specifically overlap with *foraging* gene expression in certain neuronal populations (Fig. 2E-H). This suggests that *foraging* gene expression within these memory-related brain regions is not essential for the formation of memories underlying LMD behavior, contrary to the previous assumption that such memories were primarily localized to the EB rather than the MB or FB ^15^. Therefore, the *foraging* gene may not be directly involved in the generation of LMD-related memories, or there may be other, as yet unidentified, brain regions critical for this function.

**Figure 2.**
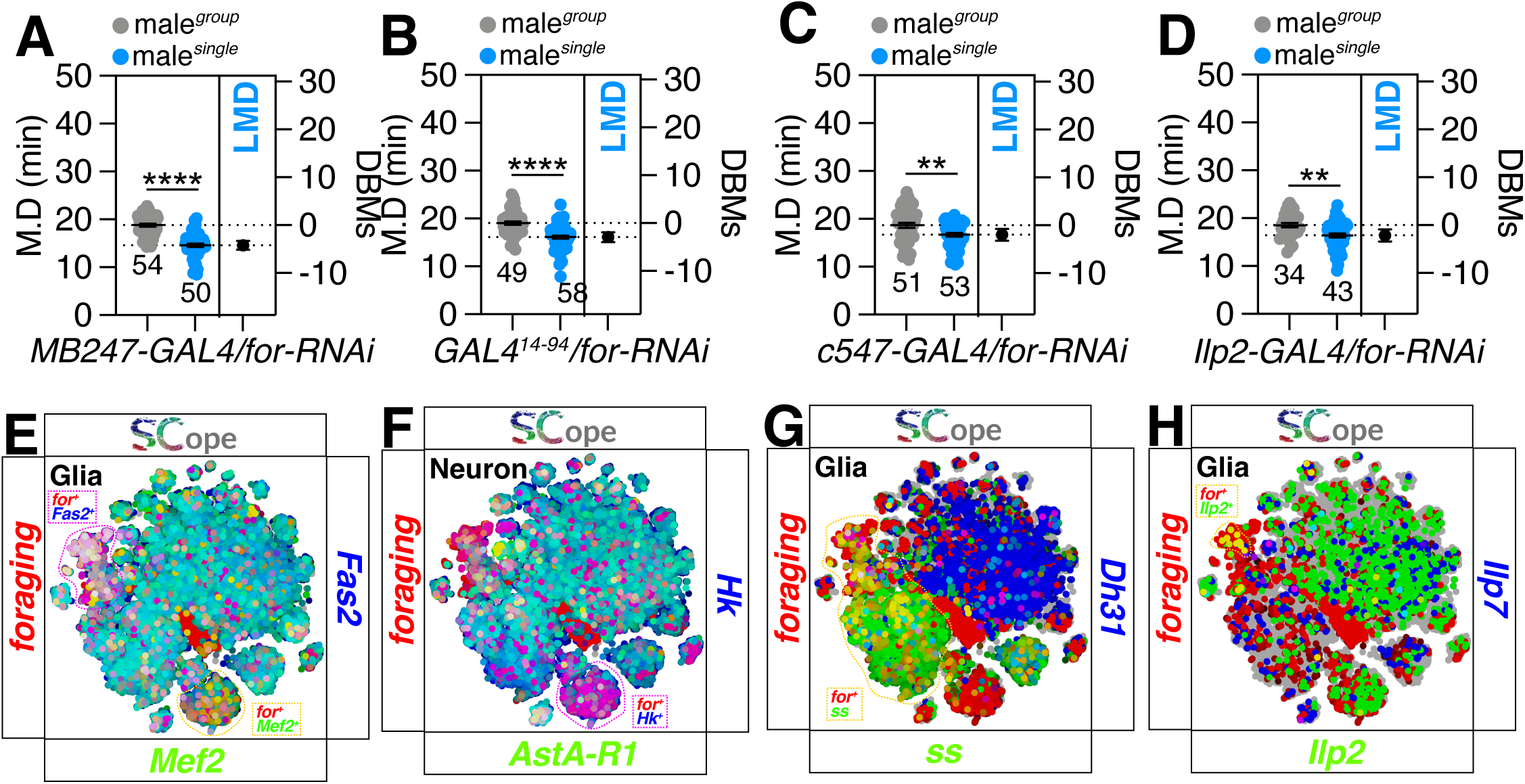
The memory circuitry responsible for LMD behavior does not correlate with *foraging* activities. (A) LMD assays of flies expressing *MB247-GAL4* (MB) driver together with *UAS-for-RNAi*. (B) LMD assays of flies expressing *GAL4^14-94^*(FB) driver together with *UAS-for-RNAi*. (C) LMD assays of flies expressing *c547*-*GAL4* (EB) driver together with *UAS-for-RNAi*. (D) LMD assays of flies expressing *Ilp2-GAL4* (PI) driver together with *UAS-for-RNAi*. (E) SCOPE scRNA-seq datasets reveal cell clusters colored by expression of *foraging* (red), *Fas2/Mef2* (blue/green) in glia cells. (F) SCOPE scRNA-seq datasets reveal cell clusters colored by expression of *foraging* (red), *Hk/AstA-R1* (blue/green) in neurons. (G) SCOPE scRNA-seq datasets reveal cell clusters colored by expression of *foraging* (red), *Dh31/ss* (blue/green) in glia cells. (H) SCOPE scRNA-seq datasets reveal cell clusters colored by expression of *foraging* (red), *Ilp7/Ilp2* (blue/green) in glia cells.

### A distinct population of peptidergic neurons that express the *for* gene are responsible for regulating LMD

LMD behavior is known to depend on the peptidergic circuitry of Pdf/NPF, but not sNPF ^16^. However, our data show that *foraging* gene expression in these peptidergic neurons does not affect LMD behavior, nor is *foraging* gene expression detected in these circuits (Fig. 3A-F). This indicates that *foraging* gene expression in the peptidergic neurons is also not required for LMD behavior associated with *foraging* gene function. Despite the absence of a clear co-expression pattern between *foraging* gene and *Pdf* or *NPF* in neuronal populations, we observed a significant overlap between the expression of *sNPF* and *foraging* gene (Fig. 3F).

**Figure 3.**
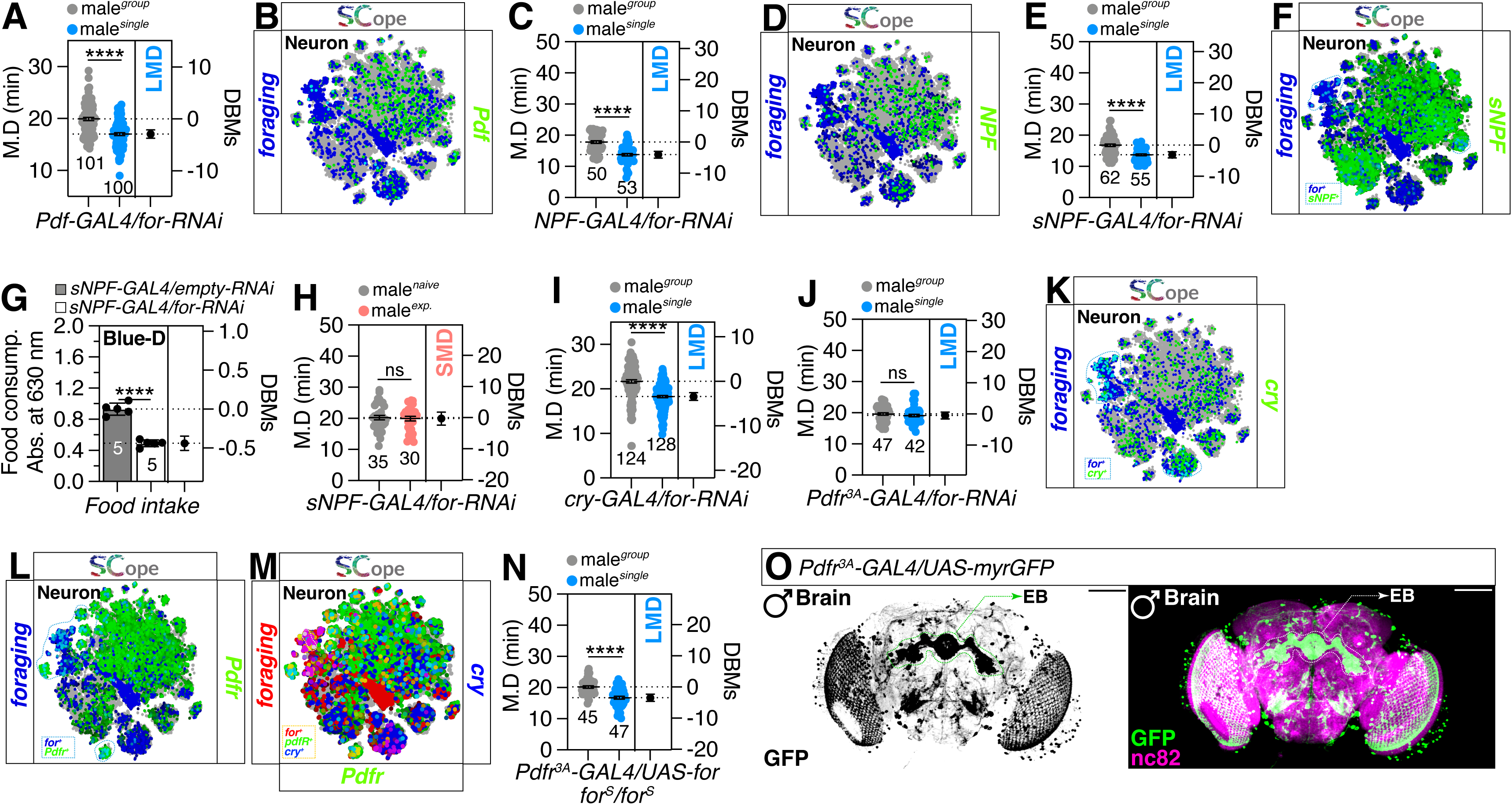
Pdfr neurons that express the *for* gene are responsible for regulating LMD. (A) LMD assays of flies expressing *Pdf-GAL4* driver together with *UAS-for-RNAi*. (B) SCOPE scRNA-seq datasets reveal cell clusters colored by expression of *foraging* (blue), *Pdf*(green) in neurons. (C) LMD assays of flies expressing *NPF-GAL4* driver together with *UAS-for-RNAi*. (D) SCOPE scRNA-seq datasets reveal cell clusters colored by expression of *foraging* (blue), *NPF* (green) in neurons. (E) LMD assays of flies expressing *sNPF-GAL4* driver together with *UAS- for-RNAi*. (F) SCOPE scRNA-seq datasets reveal cell clusters colored by expression of *foraging* (blue), *sNPF*(green) in neurons. (G) 24-h food intake of males measured by Blue-Dye assays of male flies expressing *sNPF-GAL4* driver together with *empty-RNAi/UAS-for-RNAi* (H) LMD assays of flies expressing *sNPF-GAL4* driver together with *UAS-for-RNAi*. (I) LMD assays of flies expressing *cry-GAL4* driver together with *UAS-for-RNAi*. (J) LMD assays of flies expressing *Pdfr^3A^-GAL4* driver together with *UAS-for-RNAi*. (K) SCOPE scRNA-seq datasets reveal cell clusters colored by expression of *foraging* (blue), *sNPF*(green) in neurons. (L) SCOPE scRNA-seq datasets reveal cell clusters colored by expression of *foraging* (blue), *Pdfr*(green) in neurons. (M) SCOPE scRNA-seq datasets reveal cell clusters colored by expression of *foraging* (red), *cry/Pdfr* (green/green) in neurons. (N) LMD assays for *Pdfr^3A^- GAL4* drives *for* overexpression under *for^s^* homozygote. (O) Microscopic image of male brain in which *Pdfr^3A^-GAL4* was used to simultaneously drive expression of *UAS-myrGFP* (green). Scale bars represent 100 μm.

Given that both *sNPF* and *foraging* gene are known to regulate feeding behaviors ^28,41–44^, this co-expression may account for their interconnected roles in modulating feeding-related behaviors. Consistent with our expectations, silencing the *for* gene in sNPF-expressing peptidergic neurons led to a suppression of feeding behavior (Fig. 3G). Given our recent discovery that SMD behavior relies on gustatory pathways for interval timing calculations ^19^, we postulated that *for* gene expression in these neurons might influence SMD behavior, as the circuits involved in feeding and SMD overlap. Our hypotheses were confirmed, as knocking down *for* gene in sNPF neurons led to the disruption of SMD behavior (Fig. 3H). This suggests that *for* gene is involved in gustatory modulation facilitated by sNPF-expressing peptidergic neurons.

Given that Pdf signaling is known to operate through cry- and Pdfr-positive clock neuronal circuits ^16^, we investigated the role of *foraging* gene within these circuits. We discovered that knocking down *foraging* gene expression in Pdfr-expressing cells, but not cry-expressing cells impairs LMD behavior (Fig. 3I-J), and that *foraging* gene is expressed in a specific neuronal population where *cry* and *Pdfr* are expressed (Fig. 3K-M). Furthermore, genetically restoring wild-type *foraging* gene expression in Pdfr-positive cells of the *for^S^* variant background rescues LMD behavior (Fig. 3N and Fig. S1F), indicating that Pdfr-positive cells are a crucial population for the generation of LMD behavior in the context of *foraging* gene function.

### Expression of *foraging* in Pdfr-expressing specific cell population is essential for the induction of LMD behavior

Upon examining the expression patterns of Pdfr-positive cells in the brain, we observed robust expression within the EB (Fig. 3O). Our previous research also suggested that the EB is a central region involved in forming memories related to LMD ^15^. To identify the precise region within the EB where the *foraging* gene is active in promoting LMD behavior, we employed a mini-scale screening approach using established enhancer trap GAL4 lines ^45^. These lines have been effectively utilized to delineate the anatomical structure of the adult central complex. We identified cells labeled by *30y-* and *c61-GAL4* drivers as potential candidates (Fig. 4A-K). The *30y-GAL4* driver is known for its expression in the MB, EB and FB, subesophageal ganglion (SOG), antennal & optic lobes (AL & OL), protocerebrum, and median bundle. Meanwhile, *c61-GAL4* has been associated with expression in Fa3 neurons of the central complex (CC) and extends projections through the protocerebral bridge (PB) to the FB ^45–47^. Given that the *c61-GAL4* driver, which is specific to a restricted set of neurons, disrupted LMD behavior when *for* gene expression was knocked down (Fig. 4K), while the broader EB driver *c547-GAL*4 did not (Fig. 4I), we hypothesize that a very specific subset of c61-positive neurons that are c547-negative are solely responsible for the neuronal function of *foraging* that leads to LMD.

**Figure 4.**
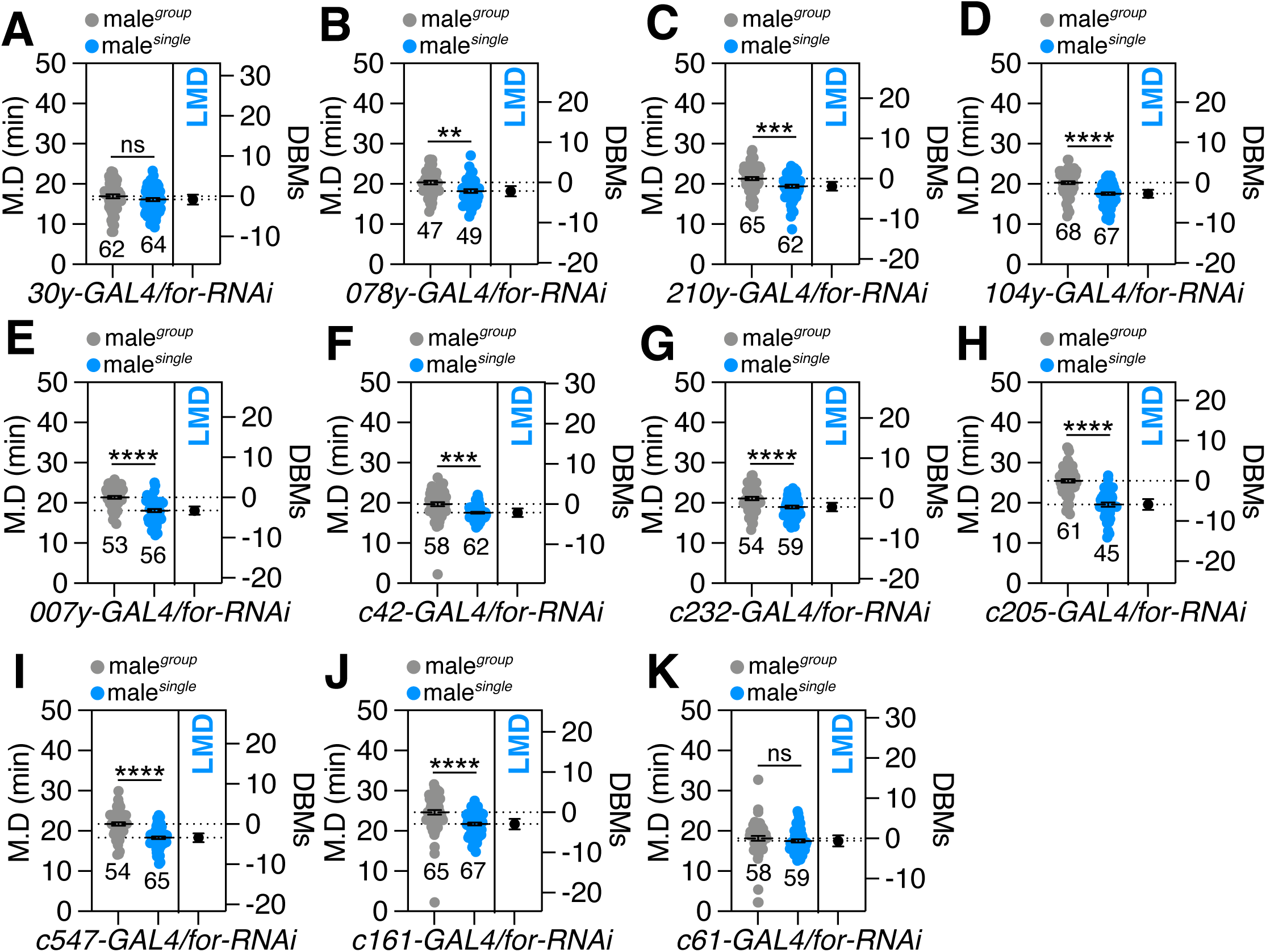
Candidate Pdfr-expressing cells in the ellipsoid body drivers for LMD screening. (A-K) LMD assays for male knockdown of *foraging* driven by subsets of neuronal cell lines of the EB,*30y-GAL4*(A)*, 078y-GAL4* (B)*, 210y-GAL4* (C)*, 104y-GAL4* (D)*, 007y-GAL4* (E)*, c42-GAL4* (F)*, c232-GAL4* (G)*, c205-GAL4* (H)*, c819-GAL4* (I)*, c161-GAL4* (J), and *c61-GAL4* (K).

Screening for overexpression of the *foraging* gene has provided evidence that the expression level of *foraging* in cells targeted by the *30y-GAL4* and *c61-GAL4* drivers is critical for the generation of LMD (Fig. S2A-K). This finding aligns with previous research indicating that feeding-related traits are influenced by the dosage of the *foraging* gene ^43^. By expressing wild-type *for* gene in cells labeled by the *30y-GAL4* driver, we were able to restore normal LMD behavior (Fig. S2L). Aligning with genetic control data (Fig. S2M), our results further suggest that the interval timing-related traits are also modulated by the dosage of the *foraging* gene within specific cell populations of the EB.

Although the specific population of neurons in the EB relies on the *foraging* gene for the induction of LMD, there may also be a non-neuronal contribution from the *foraging gene* to elicit LMD behavior. This is supported by our recent findings that corazonin receptor (CrzR), a gene expressed in glial cells, is a crucial component for LMD ^48^. Recent studies have indicated that the *foraging* gene is expressed across various tissues and is not confined to neuronal populations, extending its reach to glial subtypes and peripheral tissues such as the gastric and reproductive systems ^49^. Additionally, the expression of the *foraging* gene in multiple tissues and organs is known to be influenced by starvation ^10^. Utilizing the fly SCope 10X-Cross tissue platform, we discovered that the *foraging* gene is expressed at high levels in the majority of tissues (Fig. S3A-N). Notably, the *foraging* gene shows particularly strong co-expression with *Pdfr* in the fat body (Fig. S3A), muscle (Fig. S3I), and oenocytes (Fig. S3J).

Our findings implicate Pdf-Pdfr mediated peptidergic signaling as a mechanism through which this specific organ system influences a diverse array of behaviors and physiological processes. Considering the critical role of these organs in sugar and lipid metabolism, essential for energy storage and homeostasis, it is plausible that *foraging* gene expression within these tissues may significantly regulate energy-related behaviors and physiological responses ^50–54^. Notably, the knockdown of *foraging* gene expression in the fat body, muscle, and oenocytes did not perturb LMD behavior (Fig. S3O-Q). These results indicate that non-neuronal *foraging* gene expression is not essential for the maintenance of interval timing behaviors.

Given the compelling evidence indicating that *foraging* function is exclusively dependent on a specific neuronal population and not on other tissues for the manifestation of LMD behavior, we sought to explore the calcium response properties of *for^MI01791-TG4.1^* neurons in flies experiencing diverse social conditions. Utilizing CaLexA, a transcription-based calcium reporter system ^55^, we observed elevated levels of CaLexA signals originating from *for*-positive neurons within the antennal lobe (AL) of group-reared male flies compared to those reared in isolation (Fig. S4A-C). In contrast, the levels of CaLexA signals within the fat body remained unchanged between socially isolated and group-reared flies (Fig. S4D-E). This differential response suggests that calcium signaling within a specific neuronal population proximal to the AL plays a pivotal role in modulating interval timing behavior.

### The generation of interval timing is contingent upon the function of *foraging* in sexual dimorphic circuits

LMD represents a male-specific interval timing behavior that relies on sexually dimorphic *NPF* neurons situated within the brain ^15,16^. In *Drosophila*, sex-biased gene expression in the brain can indeed represent sexual dimorphic features ^56^. Sexual dimorphism refers to the phenotypic differences between males and females of the same species. These differences can be influenced by genetic factors, including differential expression of genes between the sexes ^57,58^.

To understand the sexually distinct expression patterns of the *foraging* gene in the brain, we examined the lacZ reporter strain *for^06860^* (Fig. 5A). Our analysis revealed that male flies exhibit a greater extent (about 2-fold) of lacZ-labeled brain regions compared to females (Fig. 5B-C). Additionally, we observed that male *foraging* gene-expressing cells closely overlap with *elav*-expressing neuronal populations (left male panels in Fig. 5A), indicating that the *foraging* gene is expressed in neuronal cells as well.

**Figure 5.**
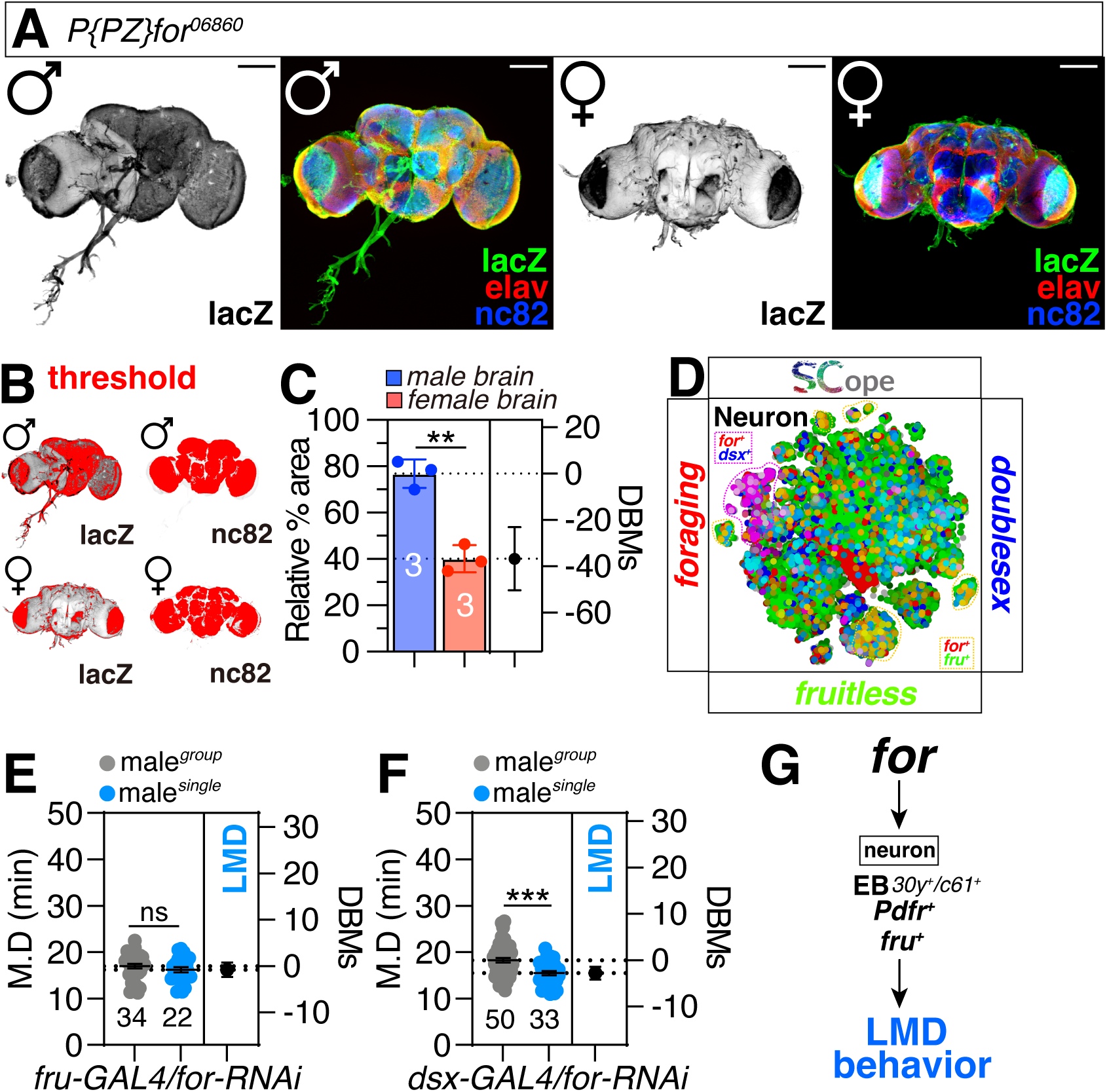
Sexually dimorphic expression of the *foraging* gene in the brain. (A) Flies expressing lacZ which is inserted in *for^06860^* were immunostained with anti-lacZ (Green), anti-elav (RFP), and nc82 (blue) antibodies. Scale bars represent 100 μm. The discrepancy in lacZ signal intensity on the surface of the male brain is likely due to the dissection procedure, which may have compromised the integrity of the surface glial cells expressing lacZ. (B) The threshold of lacZ (left panel), nc82 (right panel) in male and female were marked by threshold function of ImageJ. (C) Quantification of *foraging* gene expression in female and male brains. Fluorescence of lacZ was normalized by nc82. Bars represent the mean lacZ fluorescence level with error bars representing SEM. Asterisks represent significant differences, as revealed by the Student’s t test. (*p<0.05, **p<0.01, ***p< 0.001). (D) SCOPE scRNA-seq datasets reveal cell clusters colored by expression of *foraging* (red), *doublesex*/*fruitless* (blue/green) in neurons. (E) LMD assays for male knockdown of *foraging* driven by *fru-GAL4.* (F) LMD assays for male knockdown of *foraging* driven by *dsx-GAL4.* (G) Schematic diagram of *foraging* gene involvement in LMD.

In *Drosophila*, sexual dimorphism refers to the distinct differences in phenotype between males and females. These differences are largely determined by the expression of the *fruitless (fru)* and *doublesex (dsx)* genes, which play critical roles in the development of male and female characteristics. These genes regulate the development of male and female-specific traits, contributing to the distinct phenotypes observed between males and females of the same species^59–64^.

Investigation of the co-expression patterns of the *foraging* gene with *fru* and *dsx* revealed distinct neuronal populations that express *foraging* in conjunction with either *fru* or *dsx* (Fig. 5D). The *fru* gene is primarily involved in the regulation of sexual behaviors through the modulation of neural pathways, whereas *dsx* exhibits broader effects, impacting both neuronal circuits and the development of sexual characteristics in various cell types, extending beyond neurons ^60^. Specifically, the inhibition of the *for* gene in *fru*-expressing cells resulted in the disruption of LMD behavior (Fig. 5E), whereas the knockdown of the *for* gene in *dsx*-expressing cells did not affect LMD behavior (Fig. 5F). These findings support the hypothesis that foraging exhibits sexually dimorphic effects primarily in mal-specific neuronal populations (Fig. 5G).

Previous studies have delineated the multifaceted influence of the *foraging* gene, extending beyond locomotor behavior to include alcohol sensitivity ^7^, sucrose responses ^8^, and social network formation ^65^. These behaviors are all profoundly influenced by the serotonergic system, which plays a pivotal role in modulating a wide array of physiological and behavioral processes ^66–70^. Despite this, our experiments indicate that the *foraging* gene’s expression in the serotonergic system is not a prerequisite for LMD behavior. By manipulating *foraging* gene levels within the serotonergic system using *5-HT1B*-GAL4 and *Trhn-GAL4* drivers, we observed no disruption in LMD (Fig. S5A-D). Furthermore, we did not detect a significant co-expression between the *foraging* gene and either *5-HT1B* or *Trhn* (Fig. S5E-F), suggesting that the serotonergic system is not essential for the *foraging* gene’s function in generating LMD behavior.

## DISCUSSION

This paper investigates the influence of the *foraging* gene on interval timing behaviors in fruit flies, particularly in the context of mating duration. The *foraging* gene, which is crucial for adaptive behaviors in response to environmental cues, exhibits distinct effects in two variant phenotypes - rover and sitter ^6^. These variants present deficiencies in distinct mating duration behaviors, indicating the gene’s specific influence on interval timing (Fig. 1). Further exploration revealed the gene’s significant expression in neurons, especially those involved in memory and learning, although its impact on mating duration was not observed in key brain regions (Fig. 2). Instead, the gene’s influence was identified within specific neurons expressing Pdfr, a critical regulator of circadian rhythms, suggesting its integral role in the neural circuits governing mating duration (Fig. 3-4). Moreover, the study revealed sexually dimorphic expression patterns in the brain and the necessity of the gene’s dosage in a specific cell population for normal mating duration (Fig. 5). Collectively, these findings elucidate the complex genetic and neural circuitry underlying intricate interval timing behaviors in *Drosophila melanogaster*, highlighting the central role of the *foraging* gene (Fig. 5G).

The *foraging* gene exhibits distinct influences on interval timing behaviors (LMD and SMD) in *Drosophila*, with the rover variant specifically affecting shorter mating duration and the sitter variant affecting longer mating duration. The rescue of LMD by reintroducing wild-type *foraging* gene expression into the appropriate neurons highlights the critical role of the *foraging* gene in mediating interval timing behaviors ^8,30,43,49^. Given the utilization of distinct circadian clock genes, neuropeptides, and neural circuits by LMD and SMD ^18,22^, it would be intriguing to explore the mechanisms by which the single gene *foraging* can differentially modulate these two distinct interval timing behaviors.

The *foraging* gene’s specific role in regulating LMD was investigated through single-cell RNA sequencing and knockdown experiments ^31^. These experiments revealed the necessity of *foraging* gene expression in Pdfr-positive cells for LMD. Furthermore, the identification of key cell types in the EB that are crucial for this function when the *foraging* gene is overexpressed further supports its specific role in mediating longer mating duration. Our findings suggest the presence of a small, critical subset of EB neurons that are required to express the *foraging* gene to elicit LMD (Fig. 4 and Fig. S2). More detailed studies are required to fully understand the mechanisms underlying the generation of LMD by these specific neurons.

The sex-specific expression of the *foraging* gene in male *Drosophila* suggests its potential role in regulating male-specific behaviors. The significant overlap of *foraging* gene expression with male-specific genes *fruitless* and *doublesex* supports its involvement in male courtship and mating behaviors. However, further research is warranted to delineate the precise mechanisms by which the *foraging* gene influences sexual dimorphic behaviors.

In summary, the critical role of the *foraging* gene in mediating complex interval timing behaviors in *Drosophila* is evident. The gene’s expression in specific neurons and brain regions is crucial for its function. The intricate interplay between genetics, environment, and behavior revealed by these findings has broader implications for understanding the regulation of complex behaviors across species.

## METHODS

### Fly Stocks and Husbandry

*Drosophila melanogaster* were raised on cornmeal-yeast medium at similar densities to yield adults with similar body sizes. Flies were kept in 12 h light: 12 h dark cycles (LD) at 25°C (ZT 0 is the beginning of the light phase, ZT12 beginning of the dark phase) except for some experimental manipulation (constant dark, constant light, and experiments with the flies carrying *tub-GAL80^ts^*). Wild-type flies were *Canton-S* (*CS*). We have previously demonstrated that our *CS* flies exhibit normal LMD and SMD behaviors ^15,19^. To reduce the variation from genetic background, all flies were backcrossed for at least 3 generations to *CS* strain. Following lines used in this study, *Canton-S* (#64349), *Df(1)Exel6234*(#7708), *for^S^* (#76120), *elav^c155^* (#25750)*, repo-GAL4*(#7415)*, MB247-GAL4*(#50742) *, c547-GAL4*(#30819)*, Ilp2-GAL4*(#37516)*, Pdf-GAL4*(#6899)*, NPF-GAL4*(#25682)*, sNPF-T2A-GAL4*(#84706)*, empty-RNAi(#36304), cry-GAL4*(#24514)*, Pdfr^3A^-GAL4*(#33070)*, 30y-GAL4*(#30818)*, 078y-GAL4*(#30821)*, 104y-GAL4*(#81014)*, 007y-GAL4*(#30812)*, c42-GAL4*(#30835)*, c232-GAL4*(#30828)*, c205-GAL4*(#30826)*, c161-GAL4*(#27893)*, c61-GAL4*(#30845)*, 5-HT1B-GAL4*(#86276)*, Trhn-GAL4*(#84694)*, P{PZ}for[06860]*(#12326)*, ppl-GAL4*(#58768)*, Mhc-GAL4*(#55133)*, for^MI01791- TG4.1^ (#76140), UAS-Pdfr-RNAi*(#42508)*, UAS-myrGFP*(#77479)*, UAS-for*(#37776)*, UAS-for-RNAi*(#35158/#50741) were obtained from the Bloomington *Drosophila* Stock Center at Indiana University. *PromE(800)-Gal4[Line2M], tub-GAL80^ts^ (II)* (Korea Drosophila Resource Center,2161)*,LexAop-CD8GFP(II);UAS-CaLexA,LexAop-CD2-GFP/TM6B,Tb* (Korea Drosophila Resource Center, 1234). We thank Dr. Maria B. Sokolowski (University of Toronto) for sharing *for^R^* variants, Drs. Yuh Nung Jan and Lily Yeh Jan (UCSF, USA) for kindly sharing *GAL^14-94^* fly strain, Dr. Susan C.P. Renn (Washington University School of Medicine) for sharing *210y-GAL4* stain.

### Mating duration assay (MD assay)

The mating duration assay in this study has been reported ^15^. To enhance the efficiency of the mating duration assay, we utilized the *Df(1)^Exel6234^* (DF here after) strain in this study, which harbors a deletion of a specific genomic region that includes the sex peptide receptor (SPR) ^71,72^. Previous studies have demonstrated that virgin females of this strain exhibit increased receptivity to males ^72^. We conducted a comparative analysis between the virgin females of this line and the CS virgin females and found that both groups induced SMD. Consequently, we have selected to employ virgin females from DF in all subsequent studies. For naïve males, 40 males from the same strain were placed into a vial with food for 5 days. For single reared males, males of the same strain were collected individually and placed into vials with food for 5 days. For experienced males, 40 males from the same strain were placed into a vial with food for 4 days then 80 DF virgin females were introduced into vials for last 1 day before assay. 40 DF virgin females were collected from bottles and placed into a vial for 5 days. These females provide both sexually experienced partners and mating partners for mating duration assays. At the fifth day after eclosion, males of the appropriate strain and DF virgin females were mildly anaesthetized by CO2. After placing a single female into the mating chamber, we inserted a transparent film then placed a single male to the other side of the film in each chamber. After allowing for 1 h of recovery in the mating chamber in 25°C incubators, we removed the transparent film and recorded the mating activities. Only those males that succeeded to mate within 1 h were included for analyses. Initiation and completion of copulation were recorded with an accuracy of 10 sec, and total mating duration was calculated for each couple. We raised the flies at 22 °C from the egg stage until the adult stage for experiments that combined *tub-GAL80^ts^*. Prior to the experiment, we raised the flies at 29°C for the first 2 days to significantly stimulate the inactivation of *tub-GAL80^ts^*. We then moved them to a temperature of 25°C for the remaining 3 days to mildly induce the expression of the transgenes. The genetic controls using *GAL4/+* or *UAS/+* lines were not included in the supplementary figures because we have previously demonstrated that 100% of these flies exhibit normal LMD and SMD behaviors ^73–75^. Hence, we incorporated supplementary genetic control experiments solely if they were deemed indispensable for testing. All assays were performed from noon to 4 p.m. We conducted blinded studies for every test.

### Immunostaining and Antibodies

The dissection and immunostaining protocols for the experiments are described elsewhere ^16,19^. After 5 days of eclosion, the *Drosophila* brain has been taken from adult flies and fixed in 4% formaldehyde at room temperature for 30 minutes. The sample has been washed three times (5 minutes each) in 1% PBT and then blocked in 5% normal goat serum for 30 minutes. The sample has been next be incubated overnight at 4°C with primary antibodies in 1% PBT, followed by the addition of fluorophore-conjugated secondary antibodies for one hour at room temperature. The brain was mounted on plates with an antifade mounting solution (Solarbio) for imaging purposes. Samples were imaged with Zeiss LSM880. Primary antibodies: Chicken anti-GFP (1:500, Invitrogen), rabbit anti-LacZ antibody (1:1000, Rockland), rat anti-elav (1:100, DSHB), mouse anti-Bruchpilot (nc82) (1:50, DSHB). Fluorophore-conjugated secondary antibodies: Alexa-488 donkey anti-chicken (1:200, Jackson), Alexa Fluor 488-conjugated goat anti-rabbit (1:100, Thermo), Alexa Fluor 555-conjugated donkey anti-rat (1:100, Thermo), Alexa Fluor 647-conjugated goat anti-mouse (1:100, Jackson), plasma membranes of fatbody (stained by CellMask Deep red C10046, Thermo)

### Quantitative analysis of fluorescence intensity

To quantify the fluorescence level in brain microscopic images, we introduced ImageJ software^76^. We initially employed ImageJ’s ‘Measure’ feature to calculate average pixel intensity across the entire image or in user-specified sections, and the ‘Plot Profile’ feature to create intensity profiles across areas. To maximize precision, we converted color images to grayscale before analysis. Thresholding methods were also utilized to produce binary images that accurately outlined areas of interest, with pixel intensities of 255 (white) assigned to regions of interest and 0 (black) to the background. Intensity values from the binary image were then transferred to the corresponding locations in the original grayscale image to obtain precise intensity measurements for each object. The ‘Display Results’ feature provided comprehensive data for each object, including average intensity, size, and other relevant statistics. Fluorescence was quantified in a manually set region of interest (ROI) of brain. To compensate for differences in fluorescence between different ROI, fluorescence of lacZ and CalexA were normalized by nc82, and then the fluorescence of ROI was quantified using the measure tool of ImageJ. All specimens were imaged under identical conditions.

### Single-nucleus RNA-sequencing Analyses

The snRNAseq dataset analyzed in this paper is published in ^31^ and available at the Nextflow pipelines (VSN, https://github.com/vib-singlecell-nf), the availability of raw and processed datasets for users to explore, and the development of a crowd-annotation platform with voting, comments, and references through SCope (https://flycellatlas.org/scope), linked to an online analysis platform in ASAP (https://asap.epfl.ch/fca).

### Blue dye assay

For colorimetric food intake assay, flies were starved in PBS-containing vials for 2h and fed for 15min in vials containing 0.05% FD&C Blue dye, 7% sucrose and 5% yeast. The flies were frozen, homogenized in PBS, and centrifuged twice for 25min each. The supernatant was measured at 625nm. Each experiment consisted of 20 flies and the assay was repeated three times.

### Statistical Analysis

Statistical analysis of mating duration assay was described previously ^19^. More than 50 males (naïve, experienced and single) were used for mating duration assay. Our experience suggests that the relative mating duration differences between naïve and experienced condition and singly reared are always consistent; however, both absolute values and the magnitude of the difference in each strain can vary. So, we always include internal controls for each treatment as suggested by previous studies^77^. Therefore, statistical comparisons were made between groups that were naïvely reared, sexually experienced and singly reared within each experiment. As mating duration of males showed normal distribution (Kolmogorov-Smirnov tests, p > 0.05), we used two-sided Student’s t tests. The mean ± standard error (s.e.m) (**** = p < 0.0001, *** = p < 0.001, ** = p < 0.01, * = p < 0.05). All analysis was done in GraphPad (Prism). Individual tests and significance are detailed in figure legends. Besides traditional t-test for statistical analysis, we added estimation statistics for all MD assays and two group comparing graphs. In short, ‘estimation statistics’ is a simple framework that—while avoiding the pitfalls of significance testing—uses familiar statistical concepts: means, mean differences, and error bars. More importantly, it focuses on the effect size of one’s experiment/intervention, as opposed to significance testing^78^. In comparison to typical NHST plots, estimation graphics have the following five significant advantages such as (1) avoid false dichotomy, (2) display all observed values (3) visualize estimate precision (4) show mean difference distribution. And most importantly (5) by focusing attention on an effect size, the difference diagram encourages quantitative reasoning about the system under study^79^. Thus, we conducted a reanalysis of all of our two group data sets using both standard t tests and estimate statistics. In 2019, the Society for Neuroscience journal eNeuro instituted a policy recommending the use of estimation graphics as the preferred method for data presentation^80^.

## AUTHOR CONTRIBUTIONS

**Conceptualization:** Woo Jae Kim.

**Data curation:** Wenjing Li, Yongwen Huang,

**Formal analysis:** Wenjing Li, Yongwen Huang,

**Funding acquisition:** Woo Jae Kim.

**Investigation:** Woo Jae Kim.

**Methodology:** Woo Jae Kim.

**Project administration:** Woo Jae Kim.

**Resources:** Woo Jae Kim.

**Supervision:** Woo Jae Kim.

**Validation:** Wenjing Li, Yongwen Huang, Woo Jae Kim.

**Visualization:** Wenjing Li, Yongwen Huang, Woo Jae Kim.

**Writing – original draft:** Woo Jae Kim.

**Writing – review & editing:** Wenjing Li, Yongwen Huang, Woo Jae Kim.

## DECLARATION OF GENERATIVE AI AND AI-ASSISTED TECHNOLOGIES IN THE WRITING PROCESS

During the creation of this work, the author(s) utilized QuillBot to rephrase English sentences, verify English grammar, and detect plagiarism, as none of the authors of this paper are native English speakers. After using this tool/service, the author(s) reviewed and edited the content as needed and take(s) full responsibility for the content of the publication.

## ACKNOWLEDGEMENTS

We are very appreciative to the colleagues who supplied us with several fly strains. We thank Dr. Maria B. Sokolowski (University of Toronto) for sharing *for^R^* variants, Drs. Yuh Nung Jan and Lily Yeh Jan (UCSF, USA) for kindly sharing *GAL^14-94^* fly strain, Dr. Susan C.P. Renn (Washington University School of Medicine) for sharing *210y-GAL4* stain. This research was supported a University of Ottawa Startup grant 602496 to WJK, Startup funds from HIT Center for Life Science to WJK, a University of Ottawa Interdisciplinary Research Group Funding Opportunity (IRGFO stream 1 and 2) grants 148101 and 148747 to WJK, a Natural Sciences and Engineering Research Council of Canada (NSERC) Discovery grant (reference: 211406) to WJK, a University of Ottawa Brain and Mind Research Institute/Center for Neural Dynamics Open call project grant 150950 to WJK, a Mitacs Globalink Research Internship Program grant 17268 to WJK. This research was also supported by the Brain Pool Program of the National Research Foundation in Korea grant ZYM5041911 to WJK, Burroughs Wellcome Fund Collaborative Research Travel Grants (reference: 1017486) to WJK and a NVIDIA Academic Hardware Grant Program to WJK. The funders had no role in study design, data collection and analysis, decision to publish, or preparation of the manuscript. SGL received salary from the ‘University of Ottawa Startup grant to WJK’ and HM from the ‘Startup funds from HIT Center for Life Science to WJK’.

## SUPPLEMENTAL FIGURE TITLES AND LEGENDS

**Figure S1.**
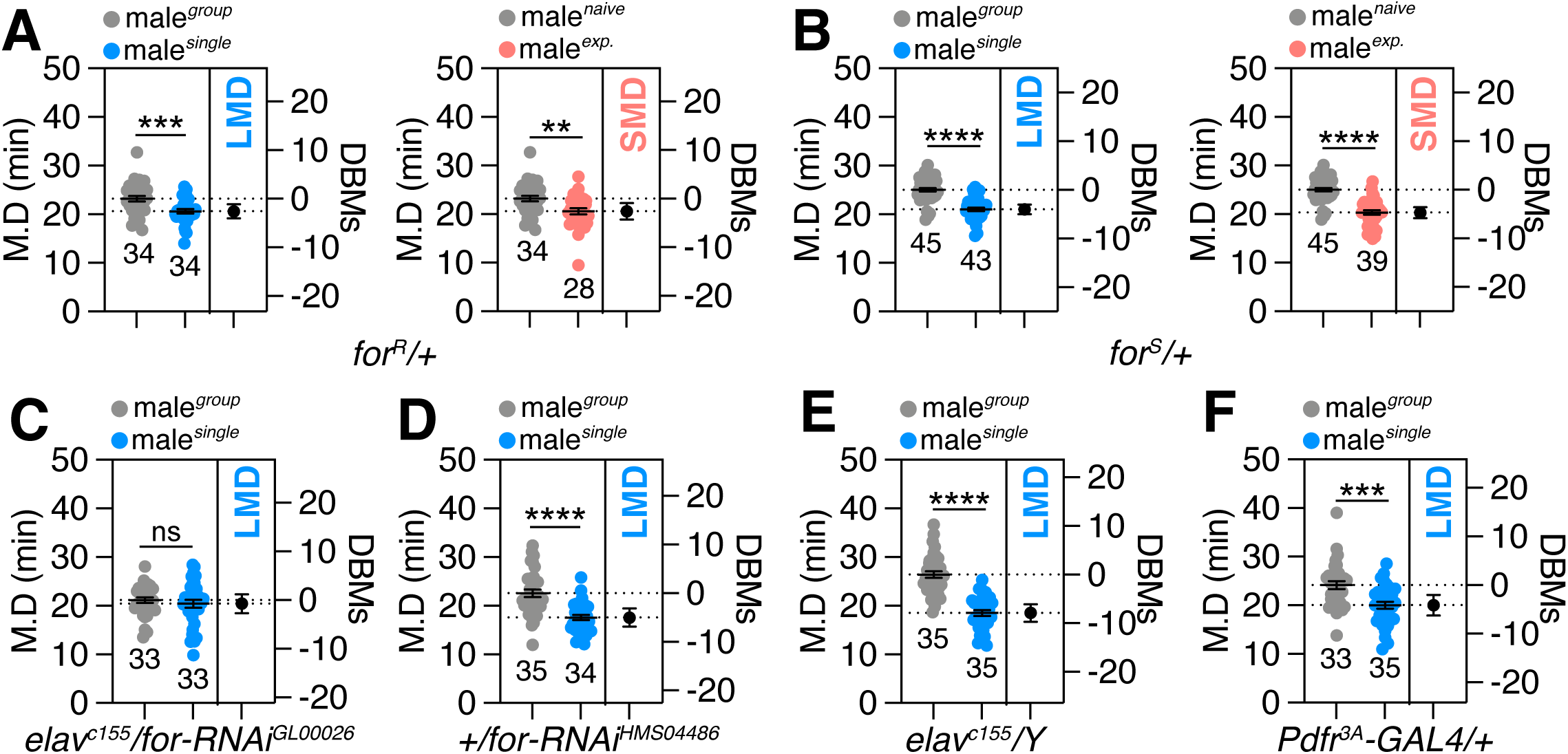
Critical Role of *foraging* Alleles in LMD and SMD behaviors. (A-B) LMD and SMD assays for *for^R^/+* and *for^S^/+* males. (C) LMD assays of flies expressing *elav^c155^* drives together with *UAS-for-RNAi^GL00026^*. (D) LMD assays for *UAS-for-RNAi^HSM04486^/+*. (E) LMD assays of flies for *elav^c155^/Y*. (F) LMD assays of flies for *Pdfr^3A^-GAL4/+*.

**Figure S2.**
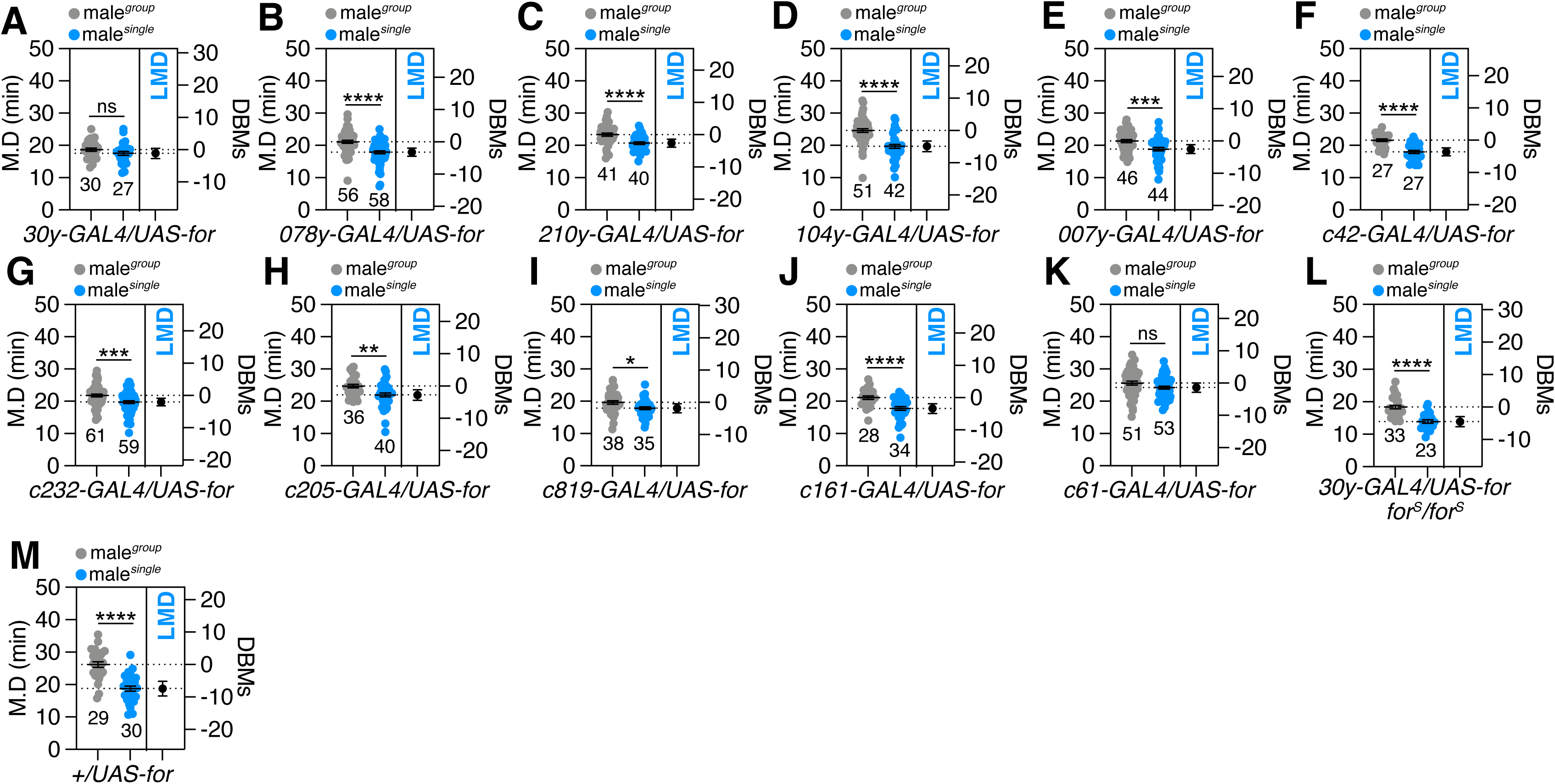
LMD Requires Specific EB Neurons with *foraging* Gene. (A) LMD assays for male overexpressing *foraging* driven by subsets of neuronal cell drivers of the EB,*30y-GAL4*(A)*, 078y-GAL4* (B)*, 210y-GAL4* (C)*, 104y-GAL4* (D)*, 007y-GAL4* (E)*, c42-GAL4* (F)*, c232-GAL4* (G)*, c205-GAL4* (H)*, c819-GAL4* (I)*, c161-GAL4* (J), and *c61-GAL4* (K). (L) LMD assays for *30y-GAL4* drives *for* overexpression under *for^S^* homozygote. (M) LMD assays for *UAS-for/+*.

**Figure S3.**
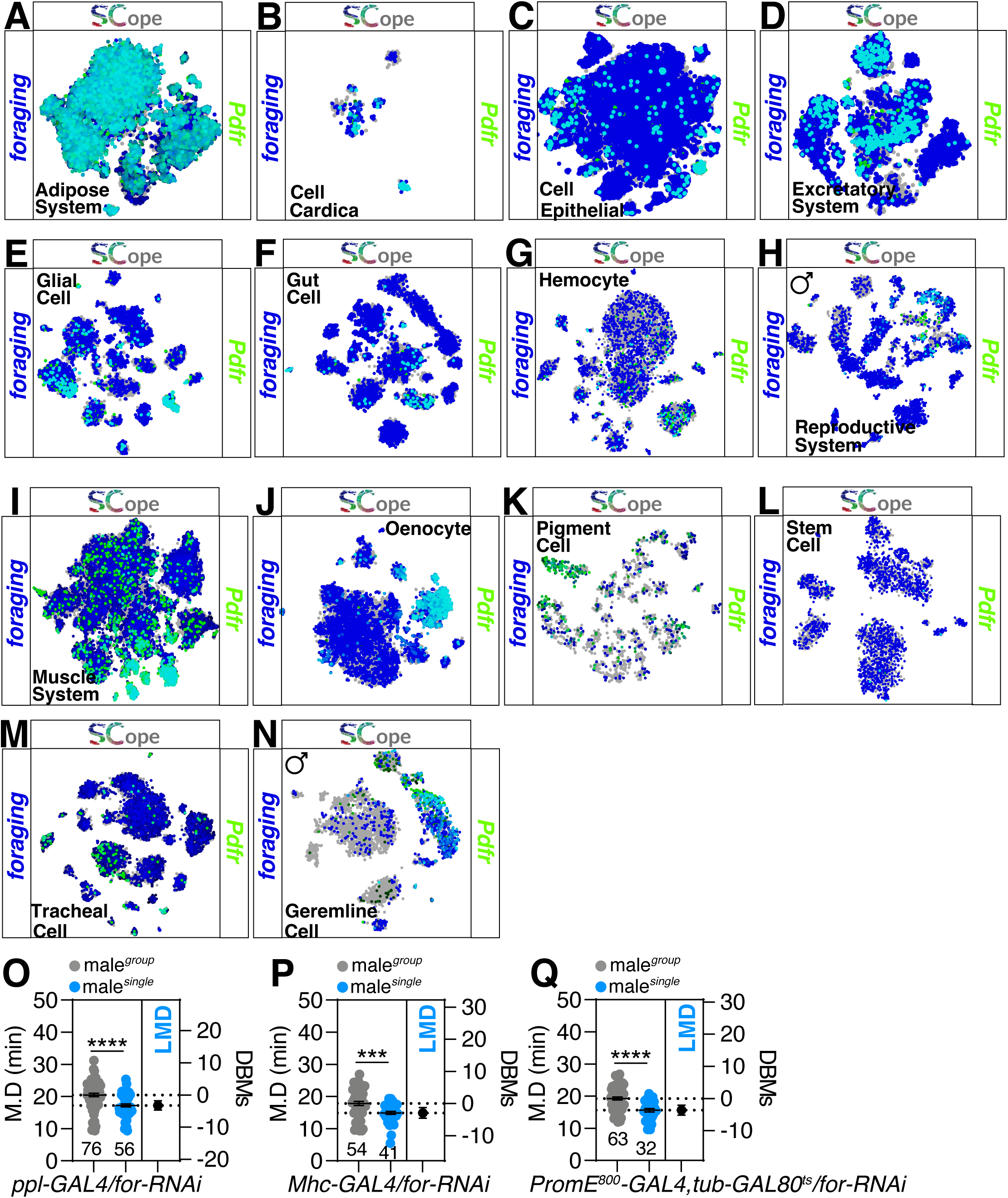
The co-expression of *foraging* gene and Pdfr in various tissues. (A-N) SCOPE scRNA-seq datasets reveal cell clusters colored by expression of *foraging* (blue), *Pdfr* (green) in different tissues. (O) LMD assays for male knockdown of *foraging* driven by *ppl-GAL4.* (P) LMD assays for male knockdown of *foraging* driven by *Mhc-GAL4.* (Q) LMD assays for male knockdown of *foraging* driven by *PromE*^800^–*GAL4* in combination with *tub-GAL80^ts^.* This allowed for the activation of *UAS-for-RNAi* expression through temperature shifts. The flies were initially raised at a temperature of 29°C for the first 2 days to substantially stimulate the expression of *UAS-for-RNAi*. They were then moved to a temperature of 25°C for the remaining 3 days to mildly induce the expression of *UAS-for-RNAi*.

**Figure S4.**
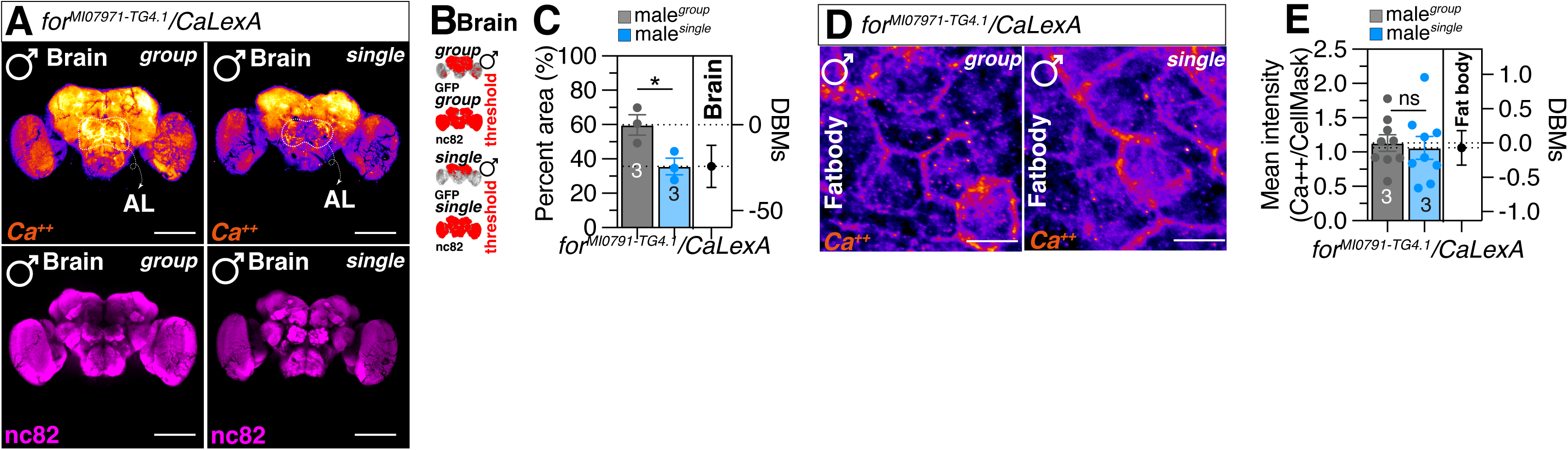
*foraging* neurons in the AL of the brain are responsible for regulating LMD. (A) Different levels of neural activity of the brain as revealed by the CaLexA system in group and single reared flies. Male flies expressing *for^MI01791-TG4.1^* along with *LexAop-CD8GFP(II);UAS-CaLexA,LexAop-CD2-GFP* were dissected after 5 days of growth. The dissected brains were then immunostained with anti-GFP (green) and anti-nc82 (blue). GFP is pseudo-colored as “red hot”. Scale bars represent 100 μm in brain panels. (B) The GFP fluorescence (green) in male fly brain was processed using ImageJ software, where a threshold function was applied to distinguish fluorescence from the background. (C) Quantification of relative value for GFP fluorescence. See the METHODS for a detailed description of the fluorescence intensity analysis used in this study. (D) Different levels of intracellular calcium level of the fat body as revealed by the CaLexA system in group and single reared flies. Scale bars represent 25 μm in fat body panels. (E) Quantification of mean intensity for GFP fluorescence. See the METHODS for a detailed description of the fluorescence intensity analysis used in this study.

**Figure S5.**
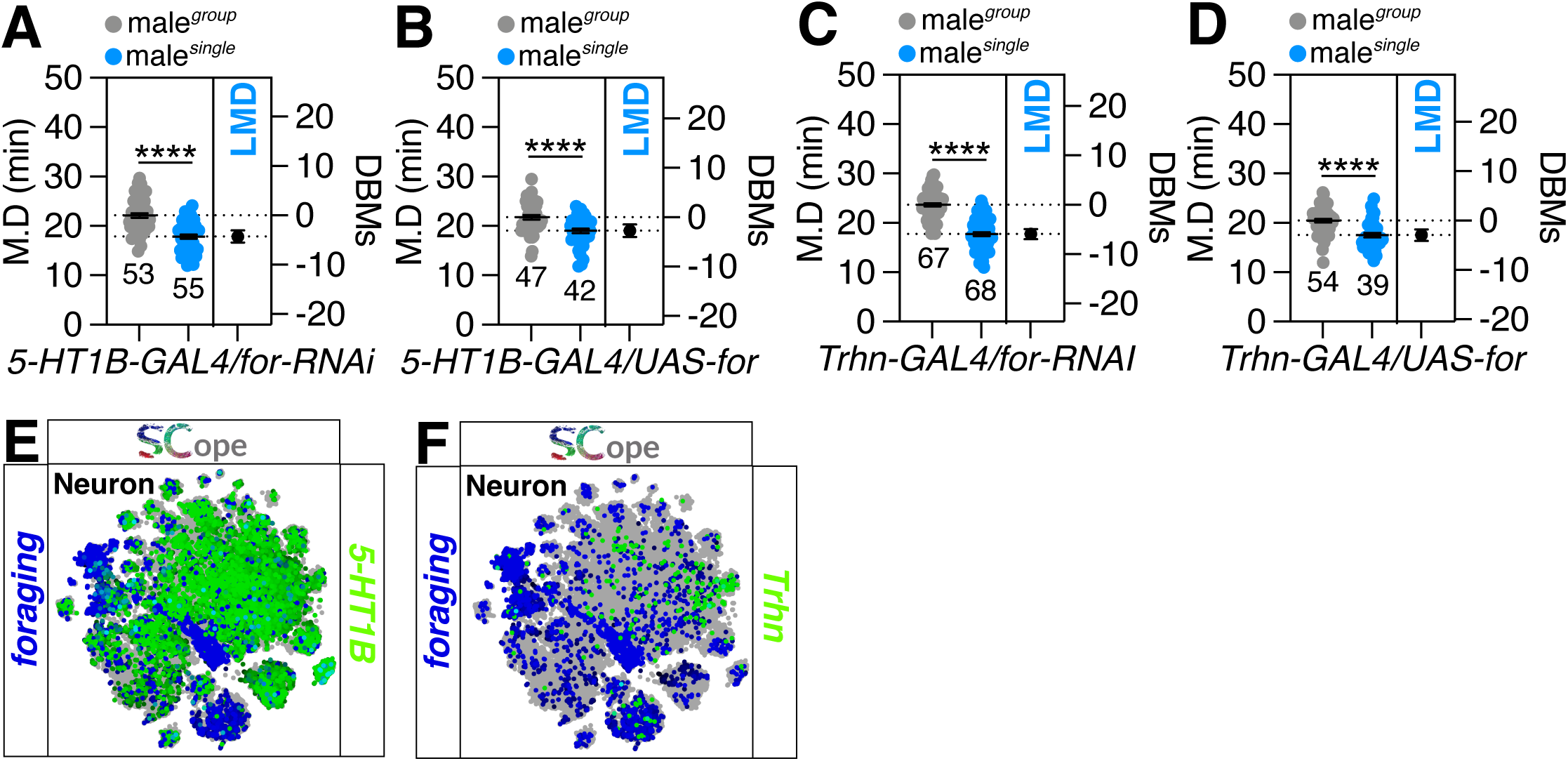
*foraging* Gene’s function Independent of Serotonergic system in LMD behavior. (A-B) LMD assays of flies expressing *5-HT1B-GAL4* drives together with *UAS-for-RNAi*(A) *and UAS-for.* (C-D) LMD assays of flies expressing *Trhn-GAL4* drives together with *UAS-for-RNAi*(C) *and UAS-for*(D). (E) SCOPE scRNA-seq datasets reveal cell clusters colored by expression of *foraging* (blue), *5-HT1B* (green) in neurons. (F) SCOPE scRNA-seq datasets reveal cell clusters colored by expression of *foraging* (blue), *Trhn* (green) in neurons.

## REFERENCES

1. Osborne, K.A., Robichon, A., Burgess, E., Butland, S., Shaw, R.A., Coulthard, A., Pereira, H.S., Greenspan, R.J., and Sokolowski, M.B. (1997). Natural Behavior Polymorphism Due to a cGMP-Dependent Protein Kinase of Drosophila. Science 277, 834–836. 10.1126/science.277.5327.834.

2. Belle, J.S. de, Hilliker, A.J., and Sokolowski, M.B. (1989). Genetic localization of foraging (for): a major gene for larval behavior in Drosophila melanogaster. Genetics 123, 157–163. 10.1093/genetics/123.1.157.

3. Ben-Shahar, Y. (2005). The foraging gene, behavioral plasticity, and honeybee division of labor. J. Comp. Physiol. A 191, 987–994. 10.1007/s00359-005-0025-1.

4. Foucaud, J., Philippe, A.-S., Moreno, C., and Mery, F. (2013). A genetic polymorphism affecting reliance on personal versus public information in a spatial learning task in Drosophila melanogaster. Proc Royal Soc B Biological Sci 280, 20130588. 10.1098/rspb.2013.0588.

5. Eddison, M., Belay, A.T., Sokolowski, M.B., and Heberlein, U. (2012). A Genetic Screen for Olfactory Habituation Mutations in Drosophila: Analysis of Novel Foraging Alleles and an Underlying Neural Circuit. Plos One 7, e51684. 10.1371/journal.pone.0051684.

6. Anreiter, I., and Sokolowski, M.B. (2019). The foraging Gene and Its Behavioral Effects: Pleiotropy and Plasticity. Annu. Rev. Genet. 53, 1–20. 10.1146/annurev-genet-112618-043536.

7. Oepen, A.S., Catalano, J., Azanchi, R., and Kaun, K.R. (2021). The foraging gene affects alcohol sensitivity, metabolism and memory in Drosophila. bioRxiv, 2021.02.09.430533. 10.1101/2021.02.09.430533.

8. Belay, A.T., Scheiner, R., So, A.K. -C., Douglas, S.J., Chakaborty-Chatterjee, M., Levine, J.D., and Sokolowski, M.B. (2007). The foraging gene of Drosophila melanogaster: Spatial-expression analysis and sucrose responsiveness. J. Comp. Neurol. 504, 570–582. 10.1002/cne.21466.

9. Wang, Z., Pan, Y., Li, W., Jiang, H., Chatzimanolis, L., Chang, J., Gong, Z., and Liu, L. (2008). Visual pattern memory requires foraging function in the central complex of Drosophila. Learn. Mem. 15, 133–142. 10.1101/lm.873008.

10. Sokolowski, D.J., Vasquez, O.E., Wilson, M.D., Sokolowski, M.B., and Anreiter, I. (2023). Transcriptomic effects of the foraging gene shed light on pathways of pleiotropy and plasticity. Ann. N. York Acad. Sci. 1526, 99–113. 10.1111/nyas.15015.

11. Buhusi, C.V., and Meck, W.H. (2005). What makes us tick? Functional and neural mechanisms of interval timing. Nat Rev Neurosci 6, 755–765. 10.1038/nrn1764.

12. Meck, W.H., Doyère, V., and Gruart, A. (2012). Interval Timing and Time-Based Decision Making. Frontiers Integr Neurosci 6, 13. 10.3389/fnint.2012.00013.

13. Boisvert, M.J., and Sherry, D.F. (2006). Interval Timing by an Invertebrate, the Bumble Bee Bombus impatiens. Curr Biol 16, 1636–1640. 10.1016/j.cub.2006.06.064.

14. Rammsayer, T.H., and Troche, S.J. (2014). Neurobiology of Interval Timing. Adv Exp Med Biol, 33–47. 10.1007/978-1-4939-1782-2_3.

15. Kim, W.J., Jan, L.Y., and Jan, Y.N. (2012). Contribution of visual and circadian neural circuits to memory for prolonged mating induced by rivals. Nat Neurosci 15, 876–883. 10.1038/nn.3104.

16. Kim, W.J., Jan, L.Y., and Jan, Y.N. (2013). A PDF/NPF Neuropeptide Signaling Circuitry of Male Drosophila melanogaster Controls Rival-Induced Prolonged Mating. Neuron 80, 1190– 1205. 10.1016/j.neuron.2013.09.034.

17. Bretman, A., Fricke, C., and Chapman, T. (2009). Plastic responses of male Drosophila melanogaster to the level of sperm competition increase male reproductive fitness. Proc Royal Soc B Biological Sci 276, 1705–1711. 10.1098/rspb.2008.1878.

18. Kim, W.J., Lee, S.G., Schweizer, J., Auge, A.-C., Jan, L.Y., and Jan, Y.N. (2016). Sexually experienced male Drosophila melanogaster uses gustatory-to-neuropeptide integrative circuits to reduce time investment for mating. Biorxiv, 088724. 10.1101/088724.

19. Lee, S.G., Sun, D., Miao, H., Wu, Z., Kang, C., Saad, B., Nguyen, K.-N.H., Guerra-Phalen, A., Bui, D., Abbas, A.-H., et al. (2023). Taste and pheromonal inputs govern the regulation of time investment for mating by sexual experience in male Drosophila melanogaster. PLOS Genet. 19, e1010753. 10.1371/journal.pgen.1010753.

20. Bretman, A., Gage, M.J.G., and Chapman, T. (2011). Quick-change artists: male plastic behavioural responses to rivals. Trends Ecol Evol 26, 467–473. 10.1016/j.tree.2011.05.002.

21. Lee, S.G., Kang, C., Saad, B., Nguyen, K.-N.H., Guerra-Phalen, A., Bui, D., Abbas, A.-H., Trinh, B., Malik, A., Zeghal, M., et al. (2022). Multisensory inputs control the regulation of time investment for mating by sexual experience in male Drosophila melanogaster. Biorxiv, 2022.09.23.509131. 10.1101/2022.09.23.509131.

22. Wong, K., Schweizer, J., Nguyen, K.-N.H., Atieh, S., and Kim, W.J. (2019). Neuropeptide relay between SIFa signaling controls the experience-dependent mating duration of male Drosophila. Biorxiv, 819045. 10.1101/819045.

23. Kim, W.J., Song, Y., Zhang, T., Zhang, X., Ryu, T.H., Wong, K.C., Wu, Z., Wei, Y., Schweizer, J., Nguyen, K.-N.H., et al. (2024). Peptidergic neurons with extensive branching orchestrate the internal states and energy balance of male Drosophila melanogaster. bioRxiv, 2024.06.04.597277. 10.1101/2024.06.04.597277.

24. Wittmann, M., and Paulus, M.P. (2008). Decision making, impulsivity and time perception. Trends Cogn. Sci. 12, 7–12. 10.1016/j.tics.2007.10.004.

25. Parent, J., Brodeur, J., and Boivin, G. (2016). Use of time in a decision-making process by a parasitoid. Ecol. Èntomol. 41, 727–732. 10.1111/een.12354.

26. Skorupski, P., and Chittka, L. (2006). Animal Cognition: An Insect’s Sense of Time? Curr. Biol. 16, R851–R853. 10.1016/j.cub.2006.08.069.

27. Thomson, J.D. (1996). Trapline foraging by bumblebees: I. Persistence of flight-path geometry. Behav. Ecol. 7, 158–164. 10.1093/beheco/7.2.158.

28. Pereira, H.S., and Sokolowski, M.B. (1993). Mutations in the larval foraging gene affect adult locomotory behavior after feeding in Drosophila melanogaster. Proc. Natl. Acad. Sci. 90, 5044–5046. 10.1073/pnas.90.11.5044.

29. Renger, J.J., Yao, W.-D., Sokolowski, M.B., and Wu, C.-F. (1999). Neuronal Polymorphism among Natural Alleles of a cGMP-Dependent Kinase Gene, foraging, in Drosophila. J. Neurosci. 19, RC28–RC28. 10.1523/jneurosci.19-19-j0002.1999.

30. Allen, A.M., Anreiter, I., Vesterberg, A., Douglas, S.J., and Sokolowski, M.B. (2018). Pleiotropy of the Drosophila melanogaster foraging gene on larval feeding-related traits. J. Neurogenet. 32, 256–266. 10.1080/01677063.2018.1500572.

31. Li, H., Janssens, J., Waegeneer, M.D., Kolluru, S.S., Davie, K., Gardeux, V., Saelens, W., David, F.P.A., Brbić, M., Spanier, K., et al. (2022). Fly Cell Atlas: A single-nucleus transcriptomic atlas of the adult fruit fly. Science 375, eabk2432. 10.1126/science.abk2432.

32. Mery, F., Belay, A.T., So, A.K.-C., Sokolowski, M.B., and Kawecki, T.J. (2007). Natural polymorphism affecting learning and memory in Drosophila. Proc. Natl. Acad. Sci. 104, 13051– 13055. 10.1073/pnas.0702923104.

33. Osborne, K.A., Belle, J.S. de, and Sokolowski, M.B. (2001). Foraging Behaviour in Drosophila Larvae: Mushroom Body Ablation. Chem. Senses 26, 223–230. 10.1093/chemse/26.2.223.

34. Crittenden, J.R., Skoulakis, E.M.C., Goldstein, E.S., and Davis, R.L. (2018). Drosophila mef2 is essential for normal mushroom body and wing development. Biol. Open 7, bio035618. 10.1242/bio.035618.

35. Kelly, S.M., Elchert, A., and Kahl, M. (2017). Dissection and Immunofluorescent Staining of Mushroom Body and Photoreceptor Neurons in Adult Drosophila melanogaster Brains. J. Vis. Exp. : JoVE, 56174. 10.3791/56174.

36. Xie, X., Tabuchi, M., Brown, M.P., Mitchell, S.P., Wu, M.N., and Kolodkin, A.L. (2017). The laminar organization of the Drosophila ellipsoid body is semaphorin-dependent and prevents the formation of ectopic synaptic connections. eLife 6, e25328. 10.7554/elife.25328.

37. Donlea, J.M., Pimentel, D., Talbot, C.B., Kempf, A., Omoto, J.J., Hartenstein, V., and Miesenböck, G. (2018). Recurrent Circuitry for Balancing Sleep Need and Sleep. Neuron 97, 378–389.e4. 10.1016/j.neuron.2017.12.016.

38. Koh, K., Joiner, W.J., Wu, M.N., Yue, Z., Smith, C.J., and Sehgal, A. (2008). Identification of SLEEPLESS, a Sleep-Promoting Factor. Science 321, 372–376. 10.1126/science.1155942.

39. Lyu, S., Terao, N., Nakashima, H., Itoh, M., and Tonoki, A. (2023). Neuropeptide diuretic hormone 31 mediates memory and sleep via distinct neural pathways in Drosophila. Neurosci. Res. 192, 11–25. 10.1016/j.neures.2023.02.003.

40. Chowański, S., Walkowiak-Nowicka, K., Winkiel, M., Marciniak, P., Urbański, A., and Pacholska-Bogalska, J. (2021). Insulin-Like Peptides and Cross-Talk With Other Factors in the Regulation of Insect Metabolism. Front. Physiol. 12, 701203. 10.3389/fphys.2021.701203.

41. Lee, K.-S., You, K.-H., Choo, J.-K., Han, Y.-M., and Yu, K. (2004). Drosophila Short Neuropeptide F Regulates Food Intake and Body Size*. J. Biol. Chem. 279, 50781–50789. 10.1074/jbc.m407842200.

42. Lee, K.-S., Kwon, O.-Y., Lee, J.H., Kwon, K., Min, K.-J., Jung, S.-A., Kim, A.-K., You, K.- H., Tatar, M., and Yu, K. (2008). Drosophila short neuropeptide F signalling regulates growth by ERK-mediated insulin signalling. Nat Cell Biol 10, 468–475. 10.1038/ncb1710.

43. Allen, A.M., Anreiter, I., Neville, M.C., and Sokolowski, M.B. (2017). Feeding-Related Traits Are Affected by Dosage of the foraging Gene in Drosophila melanogaster. Genetics 205, 761–773. 10.1534/genetics.116.197939.

44. Donlea, J., Leahy, A., Thimgan, M.S., Suzuki, Y., Hughson, B.N., Sokolowski, M.B., and Shaw, P.J. (2012). foraging alters resilience/vulnerability to sleep disruption and starvation in Drosophila. Proc. Natl. Acad. Sci. 109, 2613–2618. 10.1073/pnas.1112623109.

45. Young, J.M., and Armstrong, J.D. (2010). Structure of the adult central complex in Drosophila: Organization of distinct neuronal subsets. J. Comp. Neurol. 518, 1500–1524. 10.1002/cne.22284.

46. Tamberg, L., Jaago, M., Säälik, K., Sirp, A., Tuvikene, J., Shubina, A., Kiir, C.S., Nurm, K., Sepp, M., Timmusk, T., et al. (2020). Daughterless, the Drosophila orthologue of TCF4, is required for associative learning and maintenance of the synaptic proteome. Dis. Model. Mech. 13, dmm042747. 10.1242/dmm.042747.

47. Pauls, D., Selcho, M., Gendre, N., Stocker, R.F., and Thum, A.S. (2010). Drosophila larvae establish appetitive olfactory memories via mushroom body neurons of embryonic origin. J. Neurosci. : Off. J. Soc. Neurosci. 30, 10655–10666. 10.1523/jneurosci.1281-10.2010.

48. Zhang, T., Wu, Z., Song, Y., Li, W., Sun, Y., Zhang, X., Wong, K., Schweizer, J., Nguyen, K.-N.H., Kwan, A., et al. (2024). Long-range neuropeptide relay as a central-peripheral communication mechanism for the context-dependent modulation of interval timing behaviors. bioRxiv, 2024.06.03.597273. 10.1101/2024.06.03.597273.

49. Allen, A.M., and Sokolowski, M.B. (2021). Expression of the foraging gene in adult Drosophila melanogaster. J. Neurogenet. 35, 192–212. 10.1080/01677063.2021.1941946.

50. Tellis, M.B., Kotkar, H.M., and Joshi, R.S. (2023). Regulation of trehalose metabolism in insects: from genes to the metabolite window. Glycobiology. 10.1093/glycob/cwad011.

51. Huang, K., Liu, Y., and Perrimon, N. (2022). Roles of Insect Oenocytes in Physiology and Their Relevance to Human Metabolic Diseases. Front. Insect Sci. 2, 859847. 10.3389/finsc.2022.859847.

52. Arrese, E.L., and Soulages, J.L. (2010). Insect Fat Body: Energy, Metabolism, and Regulation. Annu. Rev. Èntomol. 55, 207–225. 10.1146/annurev-ento-112408-085356.

53. Makki, R., Cinnamon, E., and Gould, A.P. (2014). The Development and Functions of Oenocytes. Annu Rev Entomol 59, 405–425. 10.1146/annurev-ento-011613-162056.

54. Canavoso, L.E., Jouni, Z.E., Karnas, K.J., Pennington, J.E., and Wells, M.A. (2001). FAT METABOLISM IN INSECTS. Annu. Rev. Nutr. 21, 23–46. 10.1146/annurev.nutr.21.1.23.

55. Masuyama, K., Zhang, Y., Rao, Y., and Wang, J.W. (2012). Mapping Neural Circuits with Activity-Dependent Nuclear Import of a Transcription Factor. Journal of Neurogenetics 26, 89–102. 10.3109/01677063.2011.642910.

56. Khodursky, S., Svetec, N., Durkin, S.M., and Zhao, L. (2020). The evolution of sex-biased gene expression in the Drosophila brain. Genome Res. 30, 874–884. 10.1101/gr.259069.119.

57. Singh, A., and Agrawal, A.F. (2023). Two Forms of Sexual Dimorphism in Gene Expression in Drosophila melanogaster: Their Coincidence and Evolutionary Genetics. Mol. Biol. Evol. 40, msad091. 10.1093/molbev/msad091.

58. Arbeitman, M.N., New, F.N., Fear, J.M., Howard, T.S., Dalton, J.E., and Graze, R.M. (2016). Sex Differences in Drosophila Somatic Gene Expression: Variation and Regulation by doublesex. G3: Genes, Genomes, Genet. 6, 1799–1808. 10.1534/g3.116.027961.

59. Ito, H., Sato, K., and Yamamoto, D. (2013). Sex-switching of the Drosophila brain by two antagonistic chromatin factors. Fly 7, 87–91. 10.4161/fly.24018.

60. Yamamoto, D., and Koganezawa, M. (2013). Genes and circuits of courtship behaviour in Drosophila males. Nat Rev Neurosci 14, 681–692. 10.1038/nrn3567.

61. Rideout, E.J., Dornan, A.J., Neville, M.C., Eadie, S., and Goodwin, S.F. (2010). Control of sexual differentiation and behavior by the doublesex gene in Drosophila melanogaster. Nat Neurosci 13, 458–466. 10.1038/nn.2515.

62. Yamamoto, D., Fujitani, K., Usui, K., Ito, H., and Nakano, Y. (1998). From behavior to development: genes for sexual behavior define the neuronal sexual switch in Drosophila. Mech Develop 73, 135–146. 10.1016/s0925-4773(98)00042-2.

63. Pan, Y., and Baker, B.S. (2014). Genetic Identification and Separation of Innate and Experience-Dependent Courtship Behaviors in Drosophila. Cell 156, 236–248. 10.1016/j.cell.2013.11.041.

64. Pan, Y., and Baker, B.S. (2014). Genetic Identification and Separation of Innate and Experience-Dependent Courtship Behaviors in Drosophila. Cell 156, 236–248. 10.1016/j.cell.2013.11.041.

65. Alwash, N., Allen, A.M., Sokolowski, M.B., and Levine, J.D. (2021). The Drosophila melanogaster foraging gene affects social networks. J. Neurogenet. 35, 249–261. 10.1080/01677063.2021.1936517.

66. Kaun, K.R., Azanchi, R., Maung, Z., Hirsh, J., and Heberlein, U. (2011). A Drosophila model for alcohol reward. Nat. Neurosci. 14, 612–619. 10.1038/nn.2805.

67. Devineni, A.V., and Heberlein, U. (2013). The Evolution of Drosophila melanogaster as a Model for Alcohol Research. Annu Rev Neurosci 36, 121–138. 10.1146/annurev-neuro-062012-170256.

68. Hermanns, T., Graf-Boxhorn, S., Poeck, B., and Strauss, R. (2022). Octopamine mediates sugar relief from a chronic-stress-induced depression-like state in Drosophila. Curr. Biol. 32, 4048–4056.e3. 10.1016/j.cub.2022.07.016.

69. Sun, Y., Qiu, R., Li, X., Cheng, Y., Gao, S., Kong, F., Liu, L., and Zhu, Y. (2020). Social attraction in Drosophila is regulated by the mushroom body and serotonergic system. Nat. Commun. 11, 5350. 10.1038/s41467-020-19102-3.

70. Brenman-Suttner, D.B., Yost, R.T., Frame, A.K., Robinson, J.W., Moehring, A.J., and Simon, A.F. (2020). Social behavior and aging: A fly model. Genes, Brain Behav. 19, e12598. 10.1111/gbb.12598.

71. Parks, A.L., Cook, K.R., Belvin, M., Dompe, N.A., Fawcett, R., Huppert, K., Tan, L.R., Winter, C.G., Bogart, K.P., Deal, J.E., et al. (2004). Systematic generation of high-resolution deletion coverage of the Drosophila melanogaster genome. Nat. Genet. 36, 288–292. 10.1038/ng1312.

72. Yapici, N., Kim, Y.-J., Ribeiro, C., and Dickson, B.J. (2008). A receptor that mediates the post-mating switch in Drosophila reproductive behaviour. Nature 451, 33–37. 10.1038/nature06483.

73. Kim, W.J., Jan, L.Y., and Jan, Y.N. (2013). A PDF/NPF Neuropeptide Signaling Circuitry of Male Drosophila melanogaster Controls Rival-Induced Prolonged Mating. Neuron 80, 1190– 1205. 10.1016/j.neuron.2013.09.034.

74. Kim, W.J., Jan, L.Y., and Jan, Y.N. (2012). Contribution of visual and circadian neural circuits to memory for prolonged mating induced by rivals. Nat. Neurosci. 15, 876–883. 10.1038/nn.3104.

75. Lee, S.G., Sun, D., Miao, H., Wu, Z., Kang, C., Saad, B., Nguyen, K.-N.H., Guerra-Phalen, A., Bui, D., Abbas, A.-H., et al. (2023). Taste and pheromonal inputs govern the regulation of time investment for mating by sexual experience in male Drosophila melanogaster. PLOS Genet. 19, e1010753. 10.1371/journal.pgen.1010753.

76. Wiki., I. Image intensity Processing. https://imagej.net/imaging/image-intensity-processing.

77. Bretman, A., Westmancoat, J.D., Gage, M.J.G., and Chapman, T. (2011). Males Use Multiple, Redundant Cues to Detect Mating Rivals. Curr. Biol. 21, 617–622. 10.1016/j.cub.2011.03.008.

78. Claridge-Chang, A., and Assam, P.N. (2016). Estimation statistics should replace significance testing. Nat. Methods 13, 108–109. 10.1038/nmeth.3729.

79. Ho, J., Tumkaya, T., Aryal, S., Choi, H., and Claridge-Chang, A. (2019). Moving beyond P values: data analysis with estimation graphics. Nat. Methods 16, 565–566. 10.1038/s41592-019-0470-3.

80. Bernard, C. (2021). Estimation Statistics, One Year Later. eNeuro 8, ENEURO.0091-21.2021. 10.1523/eneuro.0091-21.2021.

